# Dynamics of Functional Network Organization Through Graph Mixture Learning

**DOI:** 10.1101/2021.05.25.445303

**Authors:** Ilaria Ricchi, Anjali Tarun, Hermina Petric Maretic, Pascal Frossard, Dimitri Van De Ville

**Author notes:** Equal contribution. Email address* (Ilaria Ricchi). **Correspondence** Correspondence and requests for materials should be addressed to Ilaria Ricchi.

## Abstract

Understanding the organizational principles of human brain activity at the systems level remains a major challenge in network neuroscience. Here, we introduce a fully data-driven approach based on graph learning to extract meaningful repeating network patterns from regionally-averaged time-courses. We use the Graph Laplacian Mixture Model (GLMM), a generative model that treats functional data as a collection of signals expressed on multiple underlying graphs. By exploiting covariance between activity of brain regions, these graphs can be learned without resorting to structural information. To validate the proposed technique, we first apply it to task fMRI with a known experimental paradigm. The probability of each graph to occur at each time-point is found to be consistent with the task timing, while the spatial patterns associated to each epoch of the task are in line with previously established activation patterns using classical regression analysis. We further on apply the technique to resting state data, which leads to extracted graphs that correspond to well-known brain functional activation patterns. The GLMM allows to learn graphs entirely from the functional activity that, in practice, turn out to reveal high degrees of similarity to the structural connectome. We compared similarity of the default mode network estimated from different task data and comparing them to each other and to structure. Using different metrics, a similar distinction between high- and low-level cognitive tasks arises.

Overall, this method allows us to infer relevant functional brain networks without the need of structural connectome information. Moreover, we find that these networks correspond better to structure compared to traditional methods.

## 1. Introduction

Functional magnetic resonance imaging (fMRI) is a unique tool to probe the functional architecture of the human brain. Specifically, spontaneous fluctuations of blood-oxygenation-level dependent (BOLD) signal have shown to be synchronised between brain regions during resting state (RS) despite the absence of task or external stimuli [9, 68, 69]. A repertoire of functional networks have been identified in healthy [7, 77, 66] and clinical populations [74, 45, 91]. These functional networks include both sensory regions and higher-level cognitive ones, such as the default-mode network (DMN), which generally shows reduced activity during an externally-oriented task [30], and becomes more engaged during internal mentation [2, 52].

The interest in understanding the intrinsic functional organization of the human brain has motivated many new methods to deal with RS. Functional connectivity (FC), conventionally measured as Pearson correlation between pairs of regionally-averaged time courses, allows to constitute a whole-brain functional connectome [13]. In its conventional form, FC is computed using the entire RS scan, but several “dynamic” extensions that acknowledge temporal fluctuations of FC have been proposed [15, 37, 60, 10]. One popular approach is the sliding-window technique [42, 22, 48], where time courses are segmented into temporal windows so that a time-dependent FC matrix can be obtained. Then, further analysis by graph metrics [8, 67], or dimensionality reduction methods such as singular value decomposition (SVD) [43], k-means clustering [1], or hierarchical clustering [89], can be applied to extract the most relevant FC patterns. The timescale of these dynamically-occurring FC patterns is limited by the temporal window length, and their spatial specificity by the nature of FC that characterizes interacting activity between brain regions. Modelling relationships between regions through graphs offers a wide range of benefits, from better interpretability of results, to a large number of existing methods and algorithms for further analysis [56, 55].

Bayesian and probabilistic methods represent good examples of extracting repeating activity patterns from whole-brain data [86, 81, 69]. In neuroimaging, these approaches have been explored with the aim of applying unsupervised learning to estimate different hidden temporal states; e.g., Hidden Markov Models [21] to perform an activation network analysis [82, 83, 84, 73, 85, 10]. While dynamic FC patterns are generated by looking at fluctuations in second order statistics (i.e., correlation), alternatively, instantaneous activity-based brain states can access shorter timescales and directly explain the empirical BOLD time courses. Several graph learning techniques [20, 51] have been recently proposed in order to infer a meaningful graph structure from data. Due to the instantaneous nature of these methods, the choice of crucial parameters such as window lengths [44] is eliminated.

General approaches for learning graphs to represent data structure started with Dong et al. [19] who focuses on graph Laplacian matrix inference, which enforces data smoothness on the inferred graph [38]. Using a more efficient solution, dictionary-based methods assume that signals can be modeled as a sparse combination of localised graph dictionary atoms [58] [76]. Multiple graph inference methods include works on time-varying graphs [39] [88], where temporal signals reflect the structure of several graphs, each of which is active in a pre-defined time period. Works on multi-layer graphs [63] [64] again have a predefined set of signals for each graph layer and make assumptions on similarities between graph layers. For example, the recent work of Gan et al. [26] imposes that the sparsity pattern on all inferred graphs should be similar through a Bayesian prior. All of these works focus either on inferring only one graph, or already have a predefined set of signals for each graph that needs to be inferred. Such a strong assumption is not realistic in RS fMRI data, where a wide array of dynamic functional networks are known to occur. Therefore, we appeal to a graph inference method, called Graph Laplacian Mixture Model (GLMM) [57], that simultaneously decompose signals into clusters and learns their corresponding graph structures.

Traditionally, if pairwise relationships between nodes are modelled as conditional dependencies, a graph can be seen as a sparse inverse covariance matrix of a multivariate Gaussian distribution [17, 24]. Such approach tends to reveal then the underlying structural connectivity (SC) that can be estimated using tractography applied to diffusion-weighted MRI (DW-MRI)—a link that can be quantified using different measures such as correlations [35], partial correlations and regularized versions [81, 36, 87, 47]. More recently, the graph signal processing perspective brings the focus to the inference of a graph Laplacian matrix [19], adding additional constraints to the Gaussian structure and incorporating the notion of data smoothness.

In this work, we propose a new framework to estimate multiple functional states, using a recently introduced Bayesian-based generative model that is the GLMM, suitable for mixed signals expressed on different graphs [57, 49]. Starting from the whole-brain atlas-based time courses assumed to be a mixture of multivariate Gaussian variables with different covariance structures, we cluster the brain activity in different states that are characterized by their graph representation. The three features extracted for each state are the mean activity pattern, the associated graph Laplacian, and the likelihood of occurrence over time. They bring respectively information on (i) networks activation patterns, as mean activity of brain regions, (ii) graph structure, and (iii) brain state dynamics. We first validate the proposed method by extracting distinct states of task fMRI data where the epochs are unknown to the method. The performance of the method is validated by comparing the extracted probability of occurrence of each state to the timing of the task paradigm. We also show that the extracted states consist of brain areas that are consistent with previously observed regions implicated in the corresponding task. We then apply the GLMM to RS fMRI data and obtain the most prevalent states governing spontaneously interacting brain areas.

This approach also allows us to revisit the relationship between brain function and underlying structure, which is one of the fundamental questions in neuroscience [28, 5, 8, 31, 75, 37, 61]. The traditional approaches to estimate structural connectivity from FC, mentioned earlier, do not account for fluctuations of brain activity and thus how the structure-function link is exploited differently over time. Therefore, we show that the estimated graph Laplacian matrices reveal indicative similarities with SC, using not only correlations, but also exploring a spectral-based dissimilarity score.

We observe that the most notable brain pattern that consistently is part of states in all tasks, as well as in RS, is the default mode network (DMN). This pattern is characterized by the activation of medial prefrontal cortex, posterior cingulate cortex, angular gyrus and deactivation of the other task-oriented brain regions [11]. Even though DMNs brain regions are similar among all the rest phases of the task fMRI and RS, they may vary in the way they connect. Different studies have tried to capture how the DMN differs in different neurological conditions [54, 70, 65, 62]. That is why we finally focus our attention to the similarities and differences of the DMN patterns estimated during the rest epochs across all task paradigms and during resting state. The question consequently arises as to how the differences in connectivity structure that give rise to various DMN graphs are related to the brain’s underlying anatomical structure.

Finally, leveraging on the dissimilarity score between the anatomical structure and estimated graphs, we were able to distinguish networks estimated from relatively low-level cognitive tasks (e.g., motor, language) to those reflecting higher-level of complexity such as in memory and emotional tasks. We demonstrate that the score of dissimilarity may be an index of the level of more cognitive tasks: the more dissimilar the score is, the more it indicates that that network belongs to a higher cognitive level task, suggesting an appropriate method of comparison with respect to classical Pearson correlations.

## 2. Materials and Methods

### 2.1. Data and Preprocessing

We use MRI data from the Human Connectome Project (HCP). The MRI acquisition protocols have been extensively described and presented elsewhere [27]. In particular, we used 50 subjects (see Supplementary Appendix A.5 for IDs) consisting of 4 sessions of RS scans (1200 volumes each, a total of 4800 frames), and 2 sessions each of task fMRI data (i.e., working memory, relational memory, social, language, emotion, and motor tasks). Functional volumes underwent the standard pre-processing steps [80]. All functional images were first realigned to the mean functional volume for each participant. The realigned volumes were registered to the structural T1 data using rigid-body registration (SPM12, https://www.fil.ion.ucl.ac.uk), and were detrended (i.e., constant, linear, quadratic) to remove signal drifts. Then, the images were smoothed using a Gaussian kernel with FWHM equal to 6mm. Finally, we used the Automated Anatomical Labeling (AAL, 90 regions) atlas that was re-sliced to native fMRI space to parcellate fMRI volumes and compute regionally averaged fMRI BOLD signals. The structural and diffusion-weighted MRI data of each subject were downloaded from the HCP and were processed using MRtrix3 (http://www.mrtrix.org/). Individual SCs were generated based on the total number of fibers connecting two regions divided by the volumes of connecting regions. The normalization is done to ensure that the strength of the connection is not biased towards the size of the bundles. The final SC matrix was obtained by averaging all SC matrices of all subjects.

### 2.2. Graph Laplacian Mixture Model

The GLMM is a generative model assuming that the observations belong to different types of signals with different underlying graph structures [57].

The graphs are unknown and are modeled by their graph Laplacian matrix; i.e., ***L*** = ***D*** − ***A***, where ***D*** is a diagonal matrix of node degrees, and ***A*** is the adjacency matrix.

The model estimation problem wants to fit the observed data to recover signal clusters, as well as the associated activation patterns and graph structures. Specifically, observed fMRI signals are grouped into clusters defined by different unknown brain activation patterns. The model recovers these clusters, characterized by graph Laplacians together with the probability of each cluster to occur. These graph Laplacians bring information on the brain activation patterns (estimating means), functional brain connectivity structure (Laplacians) and dynamics (probability of occurrence as function of time).

Formally, each of the *M* observed signals ***x***_*m*_ ∈ ℝ^*N*^ belongs to exactly one cluster *k* represented by the graph Laplacian ***L***_*k*_ ∈ ℝ^*N ×N*^ and mean ***µ***_*k*_ ∈ ℝ^*N*^. A binary latent variable ***z***_*m*_ ∈ ℝ^*K*^ has exactly one non-zero value, which denotes the cluster *k* that ***x***_*m*_ belongs to. A probability *α*_*k*_ defines a prior probability distribution of ***x***_*m*_ belonging to cluster *k*, namely *p*(*z*_*m,k*_ = 1) = *α*_*k*_, ∀*m*. Finally, the graph Laplacian ***L***_*k*_ ∈ ℝ^*N ×N*^ models smooth changes in signals on the corresponding graph. Large edge weight values in ***L***_*k*_ thus capture pairs of vertices that change their values in similar ways. These connections can be seen as partial correlations between two vertices in a certain state [17]. Under these assumptions, signals in each cluster *k* follow a Gaussian distribution determined through the graph Laplacian^1^ 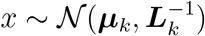 [19]:

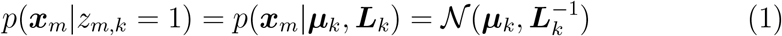

Recall that the cluster of each signal is a priori unknown. Marginalising over latent variables *z* denoting which cluster the signals belongs to, we have:

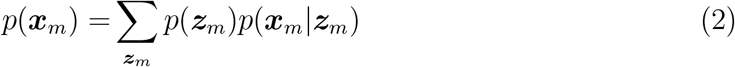

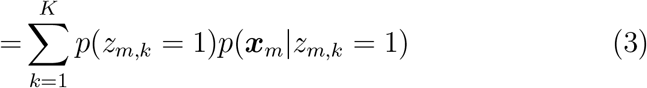

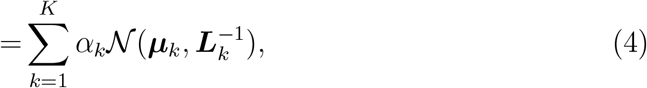

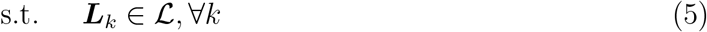

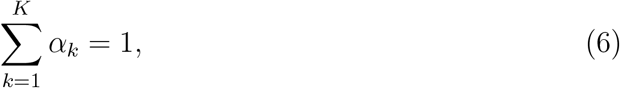

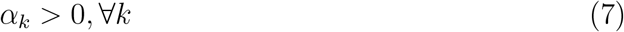

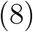

Here (5) ensures that all ***L***_*k*_’s are valid Laplacian matrices, 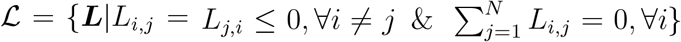. Equations (6) and (7) ensure that *α* defines a valid probability measure.

### 2.2.1. GLMM Algorithm

Given *M* observed *N* -dimensional signals in the data matrix ***X*** ∈ ℝ^*N ×M*^, we want to recover the parameters of our generative model (4). To do so, we will look at the maximum a posteriori (MAP) estimate for our parameters: probabilities ***α*** = *α*_1_, …, *α*_*K*_, means ***µ*** = ***µ***_1_, …, ***µ***_*K*_ and graph Laplacians ***L*** = ***L***_1_, …, ***L***_*K*_. Namely, we assume the data has been sampled independently from the distribution ((4)) defined through the graph Laplacians. In addition, we take into account the constraints on the graph structure given in (5), as well as possible prior information on the graphs (such as sparsity), and we maximise over the a-posteriori distribution of our model:

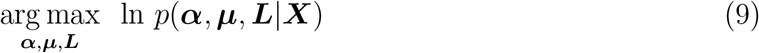

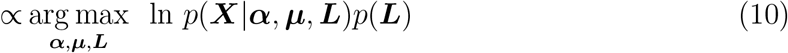

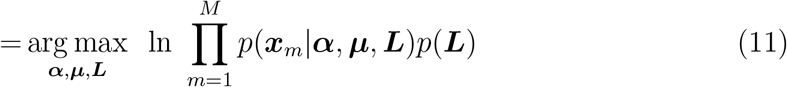

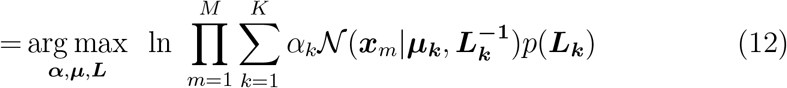

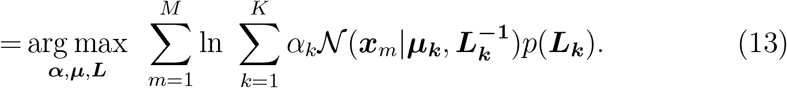

This problem does not have a closed form solution. It could be simplified through posterior probabilities ***γ*** *∈* ℝ^*M × K*^, with *γ*_*m,k*_ modeling the probability that the signal ***x*_*m*_** belongs to cluster *k*:

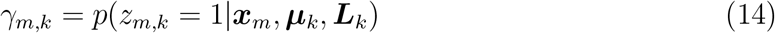

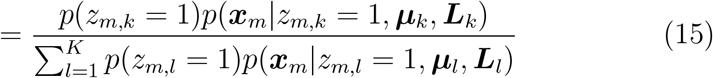

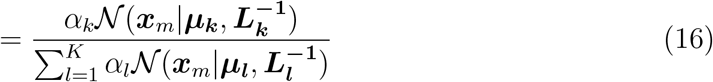

The parameters ***α, µ*** and ***L*** can now be estimated iteratively using an expectation maximisation (EM) algorithm, in which the graph Laplacians ***L*** are estimated with a graph learning scheme. For more details, we refer to the Supplementary Material [57].

### 2.3. Application to fMRI data

The general workflow of this work is summarized in Fig. 1. We build a data matrix ***X*** that contains the timecourses of all AAL 90 regions. The timecourses of all sessions and all subjects are then concatenated together to form a huge data matrix ***Y***. This is then fed to the GLMM algorithm, which learns the several graph Laplacians (***L***_*k*_), means (***µ***_*k*_) for each graph and signal clustering probabilities (***γ***).

**Figure 1:**
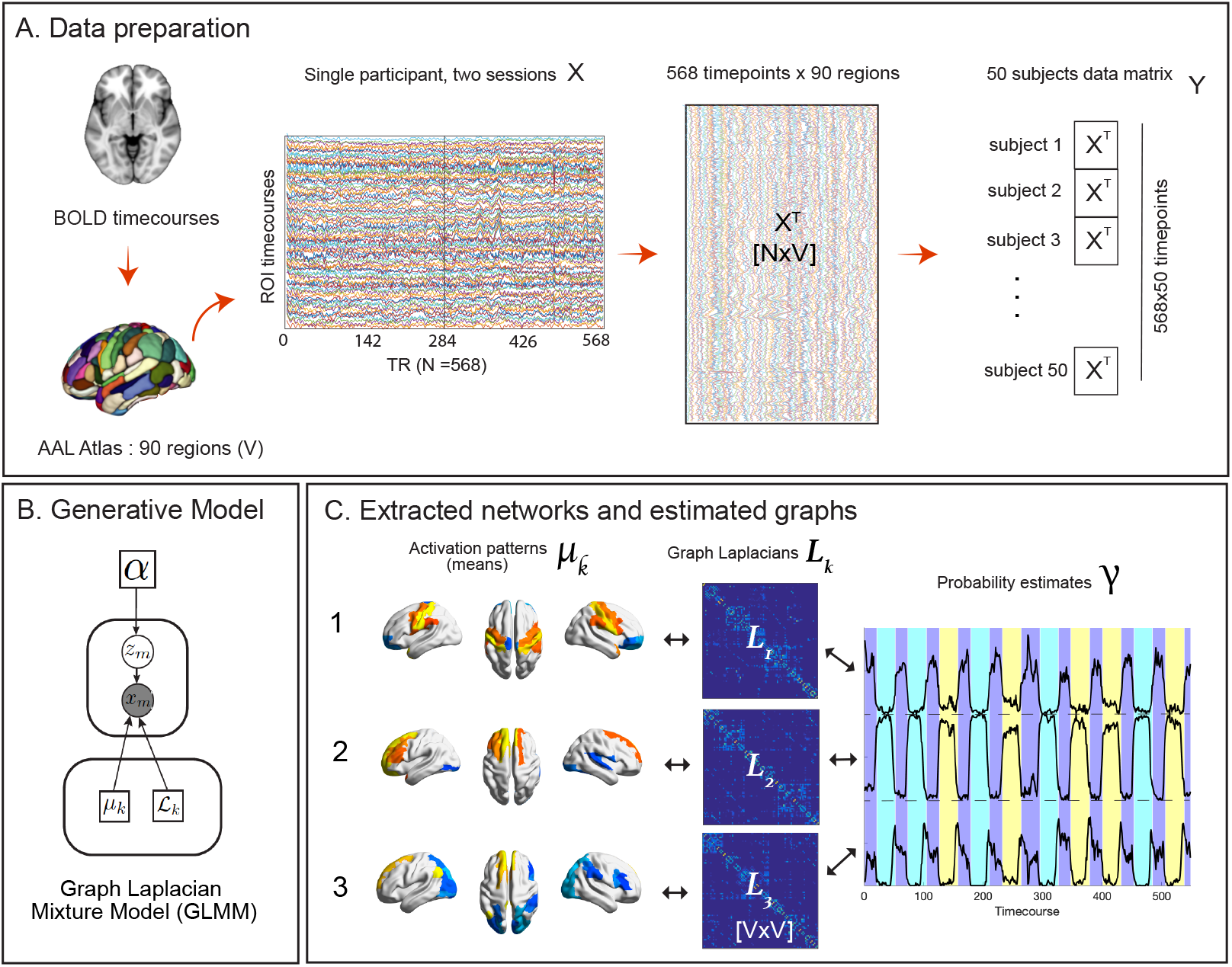
General workflow from fMRI signals to the extraction of states. (a) Mean BOLD signals from four sessions of fMRI recordings are computed within each region of the AAL90 atlas and are concatenated together to form the subject data matrix ***X***. The final data (***Y***) is obtained by concatenating 50 HCP subjects. (b) Plate notation of the generative model for extracting functional states and their corresponding estimated graphs represented through the Laplacians. (c) The output of the algorithm are *K*-number of states, their corresponding graph Laplacians, and the probability that each state would occur at a particular time-point. In the above example, we show K = 3.

We then validate the performance of the proposed framework using task fMRI as reference ground truth, by demonstrating that the timing of the task paradigms are captured by the averaged ***γ***-values across all subjects. The hyper-parameters of the model are optimized using normalized mutual information (NMI) using the task paradigm as the reference ground truth. The optimization of the number of clusters (*K*) is also guided by the consistency of the obtained clustering probabilities with the task paradigm. We cannot expect *K* to correspond to the number of tasks, since we cannot discount the possibility for a state to correspond to more than one task epoch. For that reason, we evaluate the results by iteratively changing *K* according to the concordance of the estimated ***γ*** with respect to the experimental task paradigm. In doing so, we observe that the set of meaningful network patterns is the same regardless of the proposed number of clusters *K*. Furthermore, varying *K* in such a way result in better overall accuracy, suggesting that the number of optimal clusters might be different from the number estimated through more traditional methods. We denote this method as *K*-*γ itero-homogeneity*. See Supplementary Material for further information (Fig. A2).

On the other hand, RS data lacks a ground truth, so the number of clusters *K* has been chosen based on the optimized silhouette measure and the consensus clustering procedure [53], which is a resampling-based method for optimal class discovery. *K* has eventually been varied according to the procedure mentioned above, in order to capture multiple networks. In practice, changing the optimal number of clusters does not seem to affect the final estimation, instead, it opens the possibility of inferring more networks.

### 2.4. Comparison of Learned Functional Graphs to Brain Structure

The GLMM not only solves a clustering problem but it additionally estimates a direct correlation matrix represented by the Laplacian and, differently from a Gaussian Mixture Model, the GLMM performs an implicit dimensionality reduction while leading to more interpretable results. An important benefit of the algorithm compared to other clustering methods is indeed the estimation of the *graph Laplacian*, directly estimating its inverse 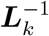. Not only does this help in obtaining more meaningful brain patterns, but it also conveys much more information and details about our clusters, allowing a direct comparison of the functional connectivity matrix with the structural connectome.

Since the DMN is strongly emerging in the state of all datasets being analyzed, they have been compared not only to each other with canonical metrics but also with SC derived from DW-MRI by using a spectral euclidean difference. This is done by first decomposing the adjacency matrices (***A***) of each graph derived from the Laplacians (***A*** = ***D*** − ***L***) into their constituent eigenvalues and eigenvectors. After a normalization step of both the functional matrix ***A*** and the structural one, the eigenspectrum of ***A*** is transformed using the Procrustes algorithm [40, 29] to match the ordering of the eigenvectors of SC.

The output of the Procrustes transform is the rotation matrix that is used to retrieve the transformed (functional) eigenvalues. In particular, given the rotation matrix ***Z*** generated by the transformation, the rotated eigenvalues are computed as follows:

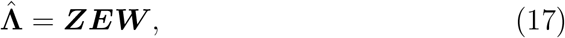

where ***W*** is the weighted adjacency matrix of the functional network taken into consideration and ***E*** is the vector of its original eigenvalues.

The spectral Euclidean difference reflects the degree of similarity between the learned graphs in each task and the anatomy defined by the SC. Thus, each network will have a certain score of distance that indicates how that function couples with structure: the smaller the metric, the closer the estimated graph to the structure. Subsequently, the scores have been normalized and sorted to see how the multiple DMNs estimated from the different tasks and RS differ from structure.

### 3. Results

### 3.1. Estimated Network time-courses are Consistent with the Timing of the Task Paradigms

The GLMM algorithm enables the recovery of three important elements: (1) means or centers of activity; (2) connectivity matrices in the form of graph Laplacians; (3) temporal activity profile of each network in terms of their likelihood to occur at each time point. The proposed framework extracts consistent patterns of brain activity for each of the considered tasks, along with a temporal profile that is strikingly in agreement with the experimental paradigm. Fig. 2 displays the estimated time-courses for six selected tasks overlaid with task conditions, which are distinguished by the background colors. States 2 and 3 of the Language task capture moments when subjects undergo the *Story* and *Math* epochs, respectively, while state 1 corresponds to the resting epoch of the task. For the *Social* task, state 3 consistently captures the RS epoch. However, the method does not distinguish the conditions *Mental* and *Random* since the plotted likelihood (***γ***) of state 2 corresponds to both task epochs. On the other hand, State 3 captures the transition between the rest and task epochs. The same observation can be made in the other tasks, in particular, the *motor, relational, and working memory*. In each of these tasks, some states capture transitions between epochs and uniquely identifies whether it is a transition between rest and task condition or a transition between two different task conditions; i.e, state 4 in *Relational Memory* captures transitions between the conditions *Relation* and *Match*, while state 2 captures transitions between rest and any of the two tasks.

**Figure 2:**
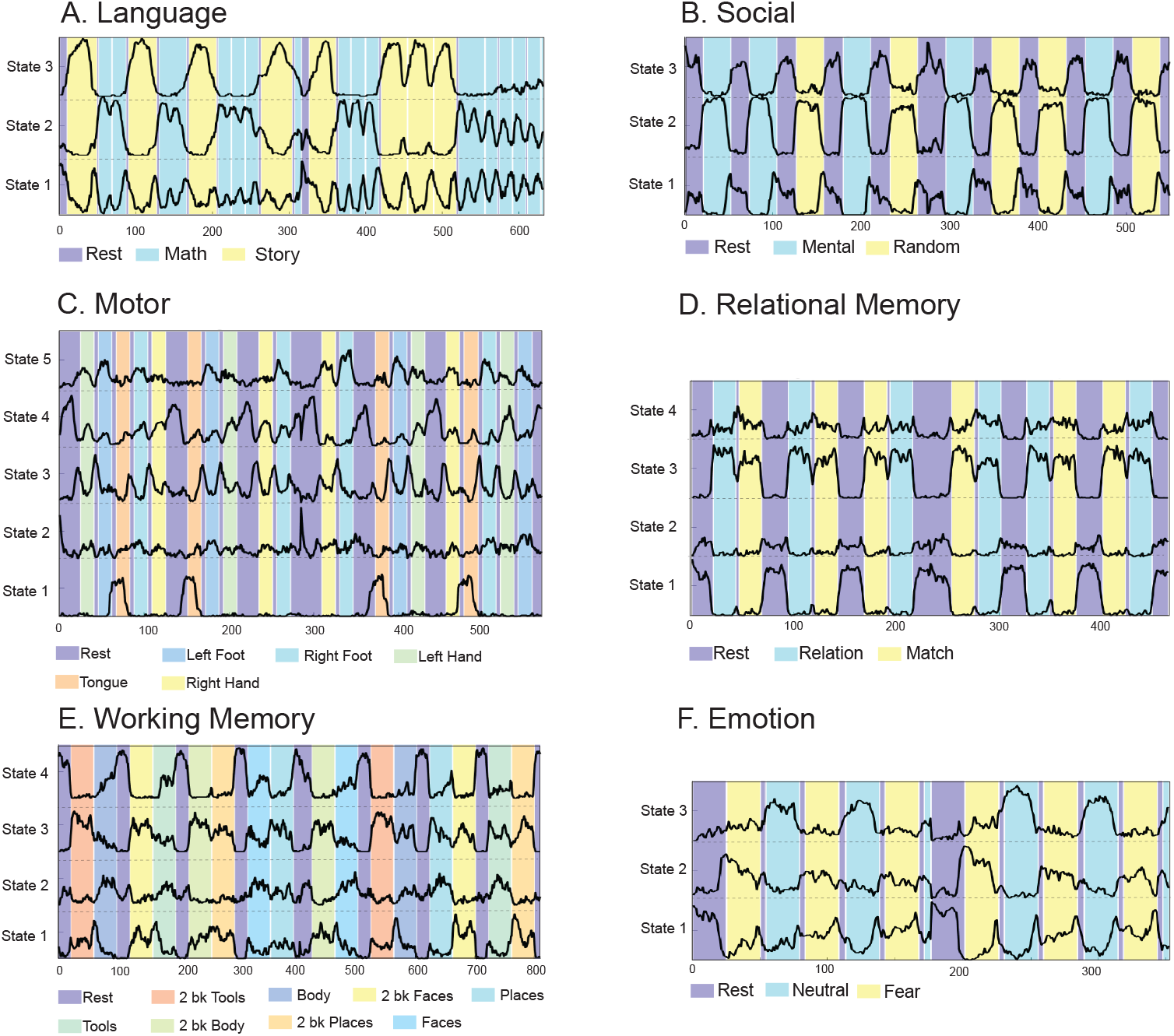
Estimated timecourses with respect to selected task paradigms. The ***γ*** values are plotted over the experimental paradigm of Language (A), Emotion (B), Social (C) and Relational Memory (D) tasks. The black signal corresponds to the probability of belonging to a specific *state*.

### 3.2. GLMM Captures Activation Patterns Corresponding To Each Task and Consistent Meta-Analytic Characteristics

Figs. 3(A) and (B) display a representative set of GLMM results for the Language and Emotion tasks, respectively. Along with the ***γ*** probabilities, already shown before, GLMM also recovers a brain graph represented with nodes whose color corresponds to the means (***µ***) estimated. Active nodes (brain areas) represented in red corresponds to more positive ***µ*** values. The states for RS epochs of both tasks reveal a spatial pattern that corresponds to the DMN network. The regions implicated in State 2 (encoding the condition *Math*) include the parietal areas (superior and inferior) and the frontal region (e.g., middle frontal gyrus, opercular part of the inferior frontal gyrus). These regions are well in-line with the established associated areas corresponding to numbers and calculations [4]. On the other hand, active areas in State 3 (condition *Story*) are the hippocampus, frontal, and the bilateral superior and anterior temporal cortex, consistent with previously observed regions implicated with story processing tasks [6].

**Figure 3:**
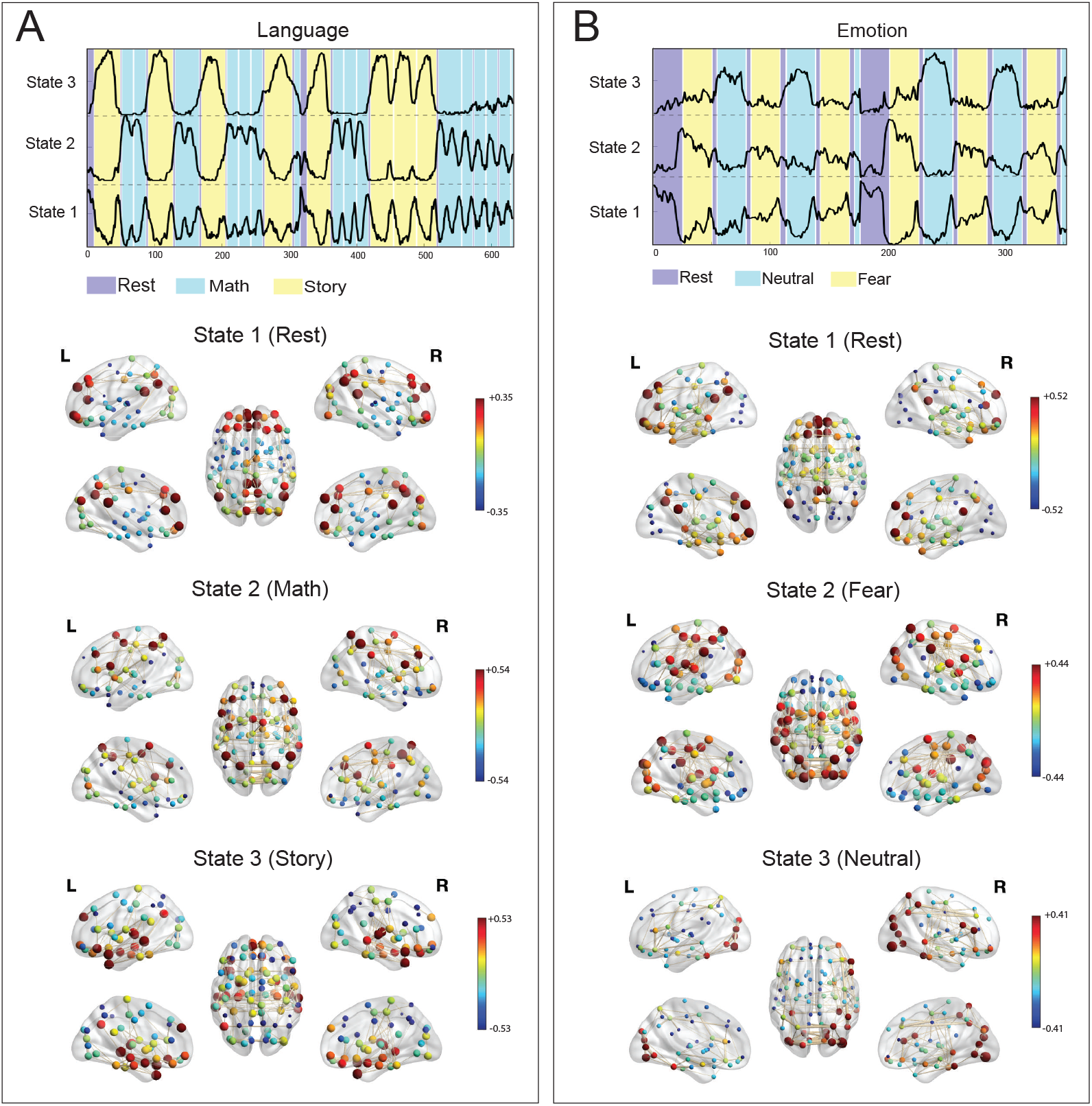
States corresponding to each epoch of the (A) Language task and (B) Emotional task paradigms. Each node corresponds to each brain region in the AAL atlas. The colors denote the values of the extracted clusters (means *µ*), the edges denote the connectivity derived from the estimated graph Laplacians, and the size the nodes indicate the degree of each node’s connections. The rest epochs of the two tasks show regions of the default mode network.

Meanwhile, states 2 and 3 of the *Emotion* task correspond to the *Fear* and *Neutral* conditions, respectively, with strong differences in terms of the regions activated. In particular, *Fear* triggers activations in the bilateral central gyrus, superior occipital gyrus, and the parietal cortices. These regions cover the somatosensory cortex, which is responsible not only for processing sensory information from various parts of the body, but also for emotional processing, including generation of emotional states and emotion regulation [18, 12, 41]. Meanwhile, the *Neutral* state shows activation of visual regions that are strongly related to shape [3, 33]. Unexpectedly, we have found relatively low means in the amygdala, a well-known region that is typically activated in emotional matching paradigms [32, 59].

### 3.3. Extracted States During Rest

After validating the outcome of the proposed framework when applied task fMRI using the known experimental task paradigm as the ground truth, we apply the method to RS fMRI. The RS brain networks that GLMM is capable of capturing are consistent with literature [79]. Fig. 4 (A) displays the estimated brain networks together with the average likelihood to occur across subjects with its standard deviation.

**Figure 4:**
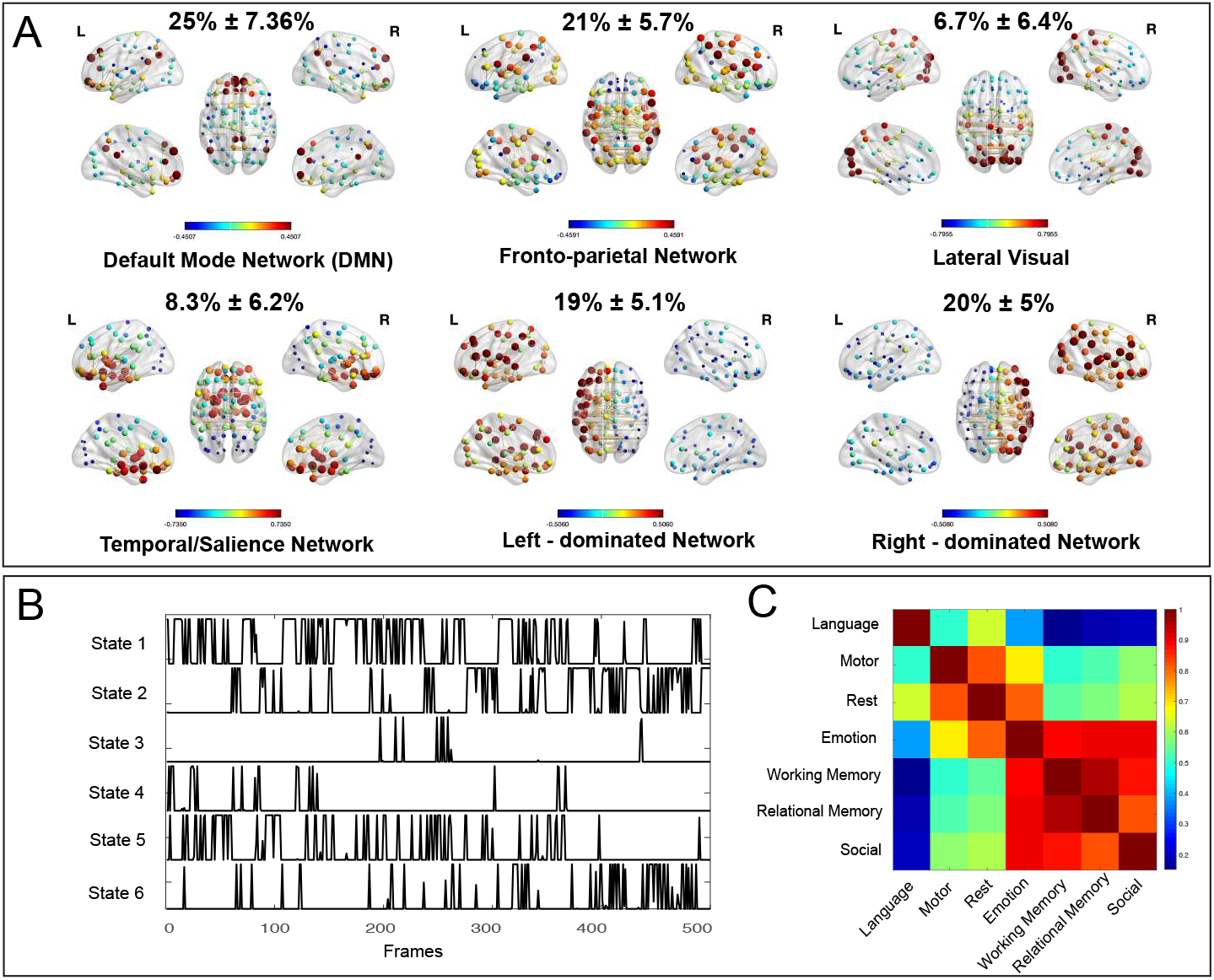
Extracted states from resting-state. (A) Six states corresponding to the RS data. Values in percentage show the mean occurrences of each state with its standard deviaition across all 50 subjects considered. (B) Example activitity profile of each state for an example subject. (C) Spatial correlation of estimated means corresponding to DMN extracted from rest and all task fMRI data.

Fig. 4 (B) shows the probability (***γ***) of each state to occur at each time-point for one representative subject, giving a proxy of the state’s activity profiles. As expected, we observe the DMN to be highly occurring. We have found a state that clearly covers areas of the visual cortex, another state that shows regions corresponding to the auditory network and some portions of the frontoparietal cortex, and another one that contains the bilateral temporal cortices and the insula, which is analogous to the salience network. Additionally, we also have found activations in the left and right hemisphere to be separately clustered. The arising networks are consistent with literature [79] but instead of estimating brain activity, GLMM instead uses an existing pool of empirical fMRI data to approximate graphs.

### 3.4. Comparison of Learned Graphs Reminiscent of the DMN

We have found that states corresponding to the rest epochs of the different tasks always bear striking similarity to the DMN, as expected. To understand the nature of the recovered DMN-related states, we compute the spatial correlation between these DMN-related means for each task as well as RS. Fig. 4(C) displays the correlation between the extracted DMN-related states from all tasks and RS fMRI. The tasks are sorted from lowest to highest correlation per row, separating low-level sensory-motor (*e*.*g*., language, motor, rest) versus high-level cognition (*e*.*g*., emotion, working memory, relational memory, and social). In particular the latters are much more similar in value.

In Supplementary A3 all the activation patterns of the DMNs estimated from all tasks and rest are shown.

### 3.5. Learned Functional DMN Graphs Comparison with Structure

Here we want to compare the DMN-related states for rest epoch of different tasks against the SC. Fig. 5(A) shows the group-averaged SC across all subjects considered in this work, and Fig. 5 (B) displays the adjacency matrix computed as ***A*** = ***D*** − ***L*** from the estimated graph Laplacian matrix of the DMN network of the Resting State. The adjacency matrix is visually more sparse than the SC. Moreover, a direct comparison (i.e., Pearson correlation across connections) between the state matrices and SC reveals similarity values within the range of *r* = 0.48 − 0.60, as shown in Fig. 5(C). It also noteworthy that, compared to the conventional FC-SC relationship, where FC is obtained by correlating inter-regional BOLD time-courses using either Pearson correlation or partial correlation, the similarity with SC is much higher for GLMM-based state matrices.

**Figure 5:**
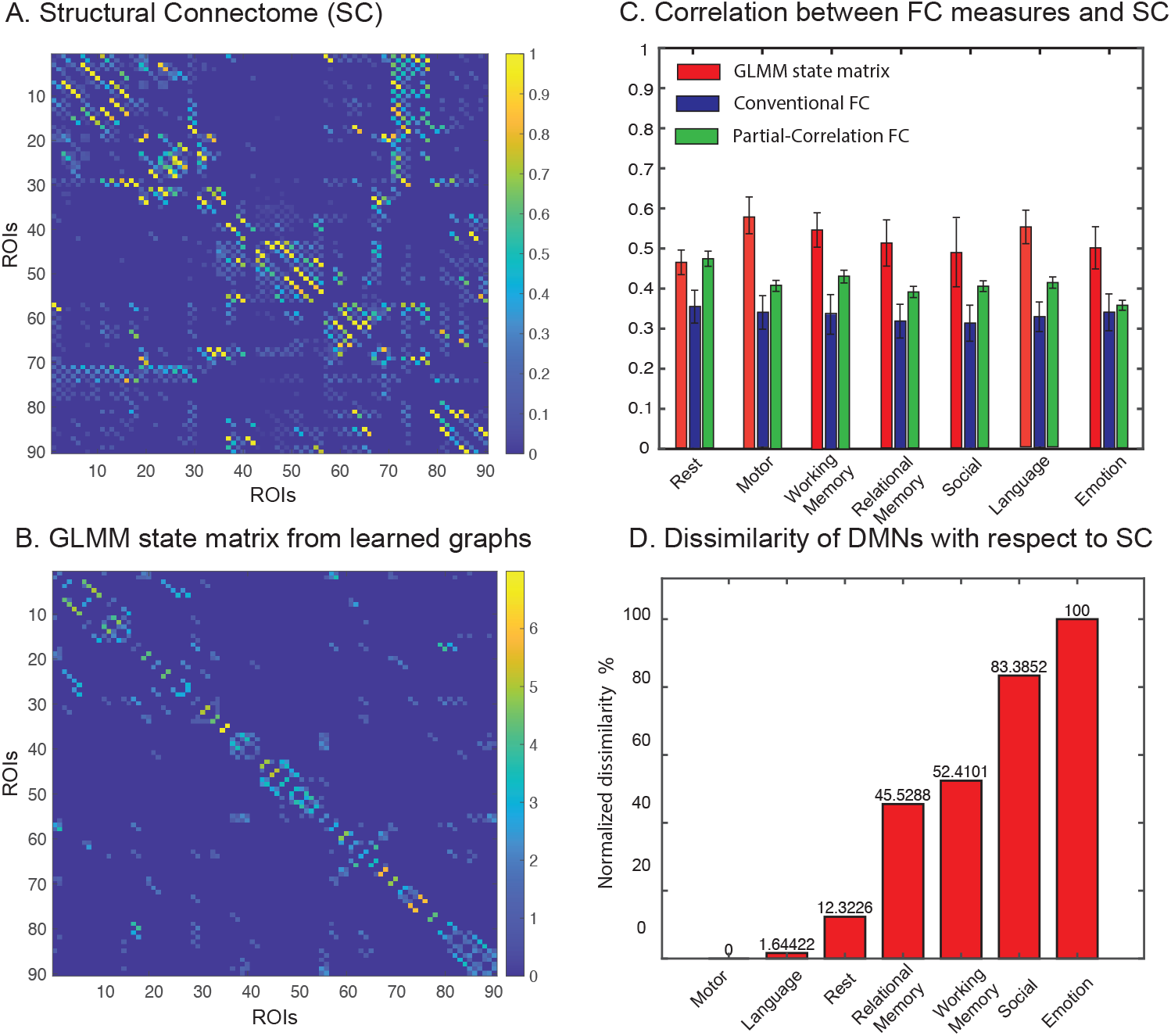
Learned Laplacian matrix and its relation to the structural connectome. (A) Group-averaged SC matrix corresponding to all subjects considered in this work. (B) Adjacency matrix extracted from the Laplacian matrix of Resting State. (C) Similarity between SC and different FC measures: using (1) GLMM-based adjacency extracted from learned graphs and classical measures of FC using (2) Pearson Correlation and (3) Partial Correlation. (D) Percentage dissimilarity scores for networks of a specific task, relative to the DMN score of that same task.

By computing the dissimilarity spectral scores for each DMN and sorting them in ascending order, it emerges that “externally directed” or less introspective tasks, such as Motor and Language, have low scores of dissimilarity with structure, hence are more similar to structure. Conversely, more introspective and “internally directed” tasks, such as Social and Emotion, differ more from structure. Thus, even though the identified brain pattern in the rest epochs across all tasks contains the key regions of the DMN, the corresponding connectivity structure of the state matrix varies, suggesting task-specific mechanisms in the way brain areas connect. Interestingly, the observed differentiation in low-to-high level cognitive tasks overlaps with the sorted correlations of pairwise DMNs previously shown in fig. 4(C).

In addition, this structure-function relation may be explored to reveal a behavioral organization of other brain patterns. In this regard, Neurosynth meta-analysis can be found in Supplementary Material (see Appendix A.1).

## 4. Discussion

### 4.1. General Findings

We have proposed the GLMM framework for explaining brain activity based on a generative model. Specifically, we were able to obtain states that are characterized by: (1) spatial pattern of regional activation levels; (2) underlying graph that describes the interactions of the regions; (3) likelihood over time.

First, we validated the approach by demonstrating that the extracted patterns are consistent with established neurophysiological descriptions corresponding to the tasks. We also showed that the probability of these states to occur at each time-point is able to capture the timing of the task paradigms. Second, we applied the approach to RS data, which revealed contributions of some of the well-known RS networks, such as the DMN, visual, auditory/attention, and salience networks. Finally, we took advantage of the estimated graph Laplacian matrices to understand the interactions of the regions, and how the strength of these interactions relates to the underlying brain structure obtained from DW-MRI. We showed that the adjacency matrices computed from the graph Laplacians bear closer similarity to SC than conventionally defined FC matrices obtained by Pearson or partial correlations of time-courses. We conclude that the graphs estimated by the GLMM are generally more correlated to structure even if the direct input is simply the empirical fMRI data, without any additional structural information.

Since a DMN-related state was found in all datasets, we have compared the spatial patterns to assess how they correlate to each other. Sorting the correlation values, a clear distinction between low- and high-level cognitive tasks was revealed. Comparison of the matrices using the spectral approach also showed a similar similar distinction. Comparing the DMN-related graphs to the SC was beneficial to evaluate how the co-activation patterns are supported in terms of connectivity, considering the estimated Laplacians that is one of the main benefits of using GLMM.

### 4.2. Estimated Graph Laplacians Conform with Structural Connectivity from DW-MRI

Pearson correlation has been widely deployed to measure FC between different breain regions [25, 60, 15, 37, 35, 72, 36]. The simplicity of the approach, however, comes with a number of well-known limitations: it only reflects a partial association between regions, with the risk of missing out time points to estimate reliable networks and it does not provide evidence of a direct relationship between two variables without confounding effects of other variables. To address this issue, partial correlation has been shown to improve the results by regressing out the effects of other variables [16, 69, 87, 47]. The GLMM approach directly estimates a Laplacian matrix to characterize the connectivity profile of a state; e.g., the GLMM adjacency matrix in Fig. 5(B) is much sparser than the FC matrix, even sparser than the SC, because of the regularization which controls sparsity. We also show in Fig. 5(C) that the GLMM adjacency matrices bear higher similarity to SC than FC variants. Unlike classical FC, GLMM adjacency matrices include direct interactions between regions within a network and do not include indirect correlations which are dependant on other regions of interest. This phenomena occurs because the GLMM, built on Gaussian Graphical models, provably have this property [78]. Moreover, the GLMM method is a dynamic approach, whereby each GLMM matrix corresponds to particular time-points (*i*.*e*., each task epoch turns out to correspond to a specific state, and consequently to a specific matrix). It has been shown in previous work [46] that there are fluctuations in similarity between FC and SC when sliding-window analysis is applied instead of static FC.

As shown in Figure 5 (C), the GLMM state matrices have closer similarity to the SC. This could be explained by the fact that these are state-specific graphs and they reflects the inherent preferential organization oth the brain to be constrained by its underlying anatomical backbone [35, 46, 14].

### 4.3. Distance Between Functional States and Structural Connectome Reflects Differences Between Tasks

The dissimilarity score based on spectral distance can be interpreted as a function-structure decoupling index of each state. In particular, each DMN-related state is associated with a value that indicates how *far away* it is from structure. This metric of function-structure *distance* was capable of separating low-from high-level tasks. As shown in previous works [52], task engagement plays a role in the regulation of DMN in order to perform goal-directed behaviors. Even though subjects are resting, they are involved with different levels of engagement.

Generally, there has always been an interest in disentangling the role of the DMN in different tasks [23]. Here, we focus on how these DMN-related states connect functionally, and their distance with anatomy as reflected by the SC. It is interesting to notice that structure-function decoupling allows distinguishing the different tasks. Thus, as shown in [23], the DMN may play a great role in internal and external tasks through a flexible coupling with task-relevant brain areas.

### 4.4. Methodological Perspectives

Dynamic analyses of functional imaging data during rest and tasks have been going on for a decade [15]. Since their inception, several methodological advances have been introduced to probe the functional organization of the brain from a dynamical point of view. A thorough review of the existing methodological tools [37, 60] has classified existing approaches into four distinct groups of methods: (i) sliding-window correlations, (ii) frame-wise analyses, (iii) state modeling, and (iv) temporal modeling. We consider our proposed technique to be an integration of state and temporal modeling, whereby the GLMM approach provides states characterized in terms of activity and connectivity, as well as the state time course, akin to how HMM models [86, 71] are able to capture various brain states and their likelihood to occur at each time-point. Unlike HMMs, however, GLMM extracts the states directly from the averaged BOLD data within parcellated brain regions with-out a dimensionality reduction step such as PCA. The reason is the implicit dimensionality reduction that occurs in GLMM by imposing a Laplacian structure on the inferred graphs [57]. This model has been shown to out-perform standard clustering methods on high-dimensional tasks, even when intuitively there is a priori no inherent graph structure in the data, achieving better clustering accuracy. Additionally, the inferred Laplacians add a strong layer of interpretability to our findings and offer multiple possibilities for further analysis and understanding of brain networks. The extracted individual graph structures describe the interactions of regions comprising the states. Apart from the enhanced visual understanding of interactions between the brain regions, these graphs enable further comparison of different brain networks, as shown in Figures 4 and 5.

Meanwhile, the extraction of the activity time course is a relevant aspect that this method highlights: in conventional methods, a functional graph is some representation of the functional connectome, which reflects the statistical interdependency between brain signals across all brain regions (nodes). The main issue with this approach is that one needs to define time windows and the size of these windows may bias the analysis [43, 34]. Furthermore, the input to the GLMM algorithm is the concatenated raw fMRI signals, and thus the experimental paradigm is completely unknown to the algorithm. Nevertheless, the ***γ*** values (probabilities of belonging to a cluster) impressively manage not only to capture the experimental conditions of the tasks, but also to extract network activity patterns that are consistent with previously established knowledge using classical regression analyses.

A particularity of the model is the imposition of a special structure on the Gaussian, bringing several benefits. Namely, as shown in [57], the model is very robust to a high-dimensional setting, when compared to standard mixtures of Gaussians. Furthermore, the graph Laplacian matrices obtained with this method offer a high level of interpretability. Therefore, the method can be applied directly to atlas-based time courses, and does not need to resolve to any prior dimensionality reduction, providing results which clearly show direct connections between different atlases.

### 4.5. Technical Limitations

While the proposed method permits to capture more information about estimated brain networks, it still suffers from some limitations, mostly due to the oversimplification of a very complex problem. Firstly, the method makes the assumption that each fMRI signal corresponds to exactly one graph, i.e., one brain network. When fitting the model, it does return probabilities of the signal belonging to each of the states, but that is largely different from assuming that one signal actually originated as a combination of several neurological networks, a phenomena that could very well be present in practice due to temporal overlap at the hemodynamic timescale. Furthermore, the method does not specify any time constraints. Even though the experimental paradigm is completely unknown to the algorithm, the ***γ*** values (probabilities of belonging to a state) manage to capture the timing of the experimental conditions. This observation helped us to validate the method, showing that even with no temporal information, meaningful states are obtained. How-ever, in practice the temporal information is valuable and incorporating it could lead to more accurate approximation of the networks and associated graphs.

Finally, additional information from the literature could be incorporated into the graph inference problem, so that more accurate and reliable findings can be obtained. These limitations can be addressed in future work.

## 5. Conclusion

This study presents a new framework for uncovering dynamic representations of brain activity. The extracted state graphs are sparse and mainly capture connections supported by the underlying SC. We demonstrate that the degree of dissimilarity between DMN-related states and SC allows to distinguish externally and internally directed tasks; i.e., low-level sensory-motor processing being more similar to structure, and high-level cognitive processing more discordant. Overall, our findings validate the potential of the proposed technique in providing a meaningful representation of brain activity.

## Acknowledgements

This work was supported by the Swiss National Science Foundation under the Project Grant 205321-163376. We thank Dr. Maria Giulia Preti for providing guidelines on the meta-analysis with Neurosynth.

## Author Contributions

**Ilaria Ricchi**: Software, Validation, Formal analysis. Writing - Original draft - Review and Editing. **Anjali Tarun**: Conceptualization, Formal analysis, Data curation, Writing - Original draft - Review and Editing, Supervision. **Hermina Petric Maretic**: Methodology, Conceptualization, Writing - Original draft, Writing - Review and Editing, Supervision. **Pascal Frossard**: Methodology, Writing - Review and Editing, Supervision. **Dimitri Van De Ville**: Conceptualization, Writing - Review and Editing, Supervision.

### Appendix A. Supplementary material

#### Appendix A.1. Structure-function relation reveals behaviorally-relevant organization

As a potential application, we performe a meta-analysis [90] of this SC-FC relationship by using spectral difference as a measure of decoupling index between SC and FC. This choice is motivated by the SC-FC decoupling measure introduced in previous work [61]. In order to be consistent, we opt for using a dissimilarity measure instead of similarity (*e*.*g*., correlation).

Similarly to the meta-analysis implemented by Margulies *et*.*al*. [50] and Preti *et*.*al*. [61], behavioral topic terms from the Neurosynth database ^2^ are matched with our decoupling global measure.

We demonstrate that the degree of their similarity captures a behaviorally-relevant gradient that is consistent with the previously observed macro-scale organization of the cortex that ranks low-level sensory processing to high-level cognitive ones [61, 50].

We perform 100 of bootstrap runs to extract networks that show statistically different values. The active areas of the estimated networks are correlated with the database present in literature, thereby capturing a number of possible behavioral domains. The same 24 topics adopted in these two studies have been considered. This behavioral gradient has been sorted according to a weighted mean of the resulting z-statistics.

#### Appendix A.2. Hyper-parameters of the model

The hyper-parameters of the model are K, Δ and *θ*. The last two are strictly related to the graph structure following the equation 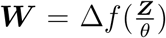, where f is the function that computes a weighted adjacency matrix ***W*** from squared pairwise distances in ***Z***, using the smoothness assumption that *trace*(***X***^***T***^ ***LX***) is small, where ***X*** is the data (columns) changing smoothly from node to node on the graph and ***L*** = ***D* − *A*** is the combinatorial graph Laplacian. See [57] for the theory.

Recall that the Gaussians depend on the inverse of the Laplacian matrix, so increasing Δ, the smaller and less overlapping the Gaussians become. The parameter *θ* represents the sparsity of the estimated adjacency matrix. By decreasing *θ*, the pairwise distance increases and the adjacency matrix becomes sparser.

**Figure A1:**
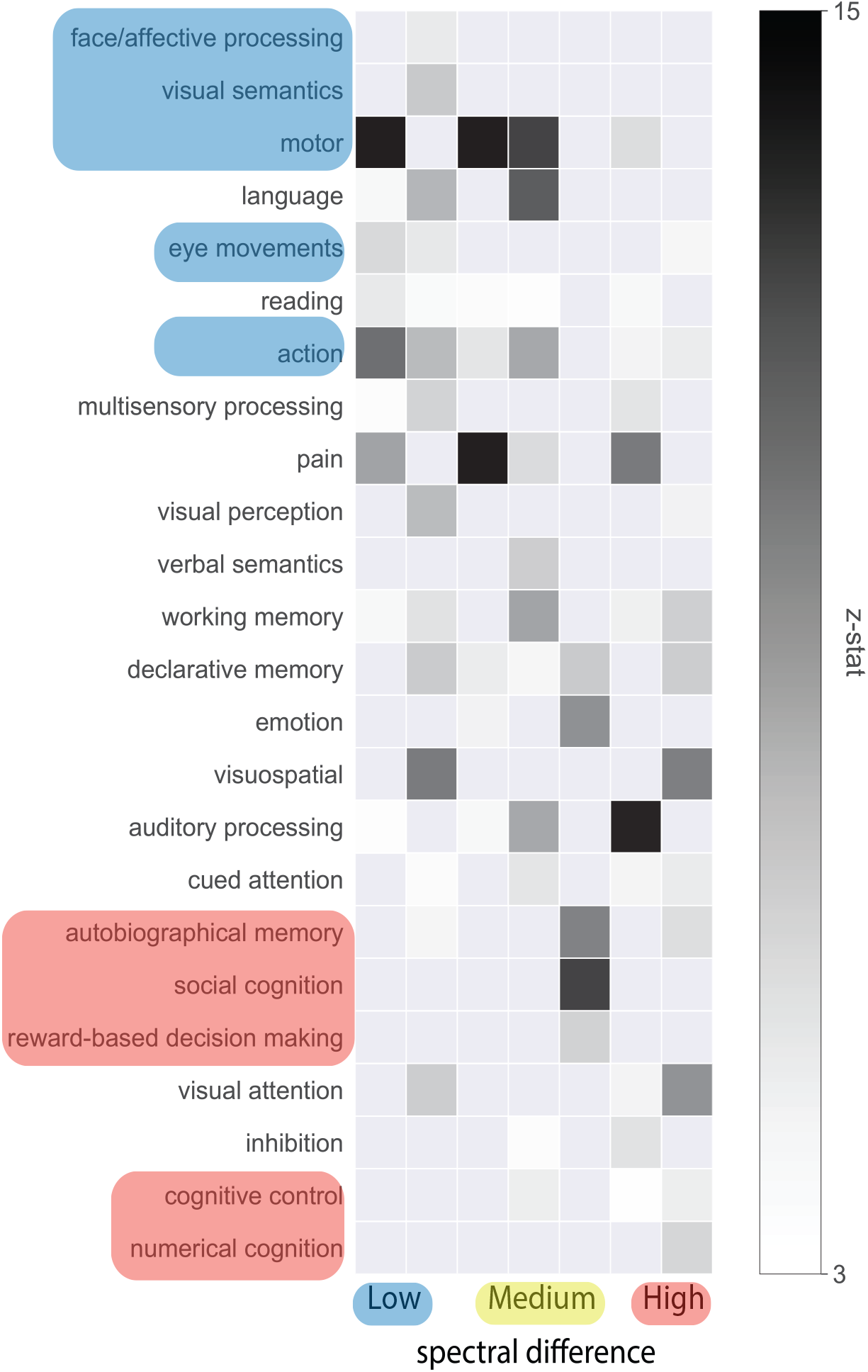
Meta-analysis of SC-FC relationship using spectral distance as a metric. The distinction between low level and high-level processing arises thanks to the sorting of the spectral difference and the z-score assigned to a behavioral topic.

All hyper-parameters of the model are optimized using a simple grid search maximizing the Normalized Mutual Information (NMI), using the task paradigm as the reference ground truth.

NMI is the metric used for task fMRI, having ground truth labels, while silhouette is used for RS.

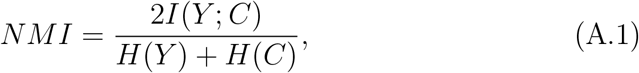

where C are the clusters labels (task paradigm), Y the class labels estimated, H is the entropy and I is the mutual information.

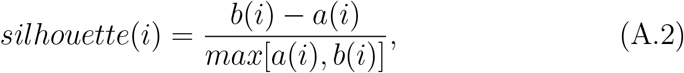

where i is the single data point, a is the average distance between i and all other data points in the same cluster and b the smallest average distance of i to all points in any other cluster.

#### Appendix A.2.1. K-γ itero-homogeneity

The change of K does not affect the estimation: the occurrence of some specific states have been compared to the experimental paradigm and in some cases changing K from the optimal generated a higher occurrence. Doing so, the ***γ*** estimation (probability of a time sample to belong to a state) gets closer to the ground-truth.

In Fig. A2 (B) ***γ*** values are plotted differently to better appreciate their values: on the x axis there are the K states, on the y axis the timecourse and the colors represent the ***γ*** values. The optimal number of cluster is supposed to be 4, but reducing K to 3 and eventually to 2 increases the probability of belonging to a cluster, in fact, the color yellow becomes brighter, showing that not necessarily the optimal K found with the NMI optimization is the *best* in capturing the dynamic.

#### Appendix A.3. Stability

The algorithm has been proven to be stable: the Hungarian algorithm has been applied to associate clusters of multiple runs and, indeed, it is able to associate the same inferred networks in the different iterations.

**Figure A2:**
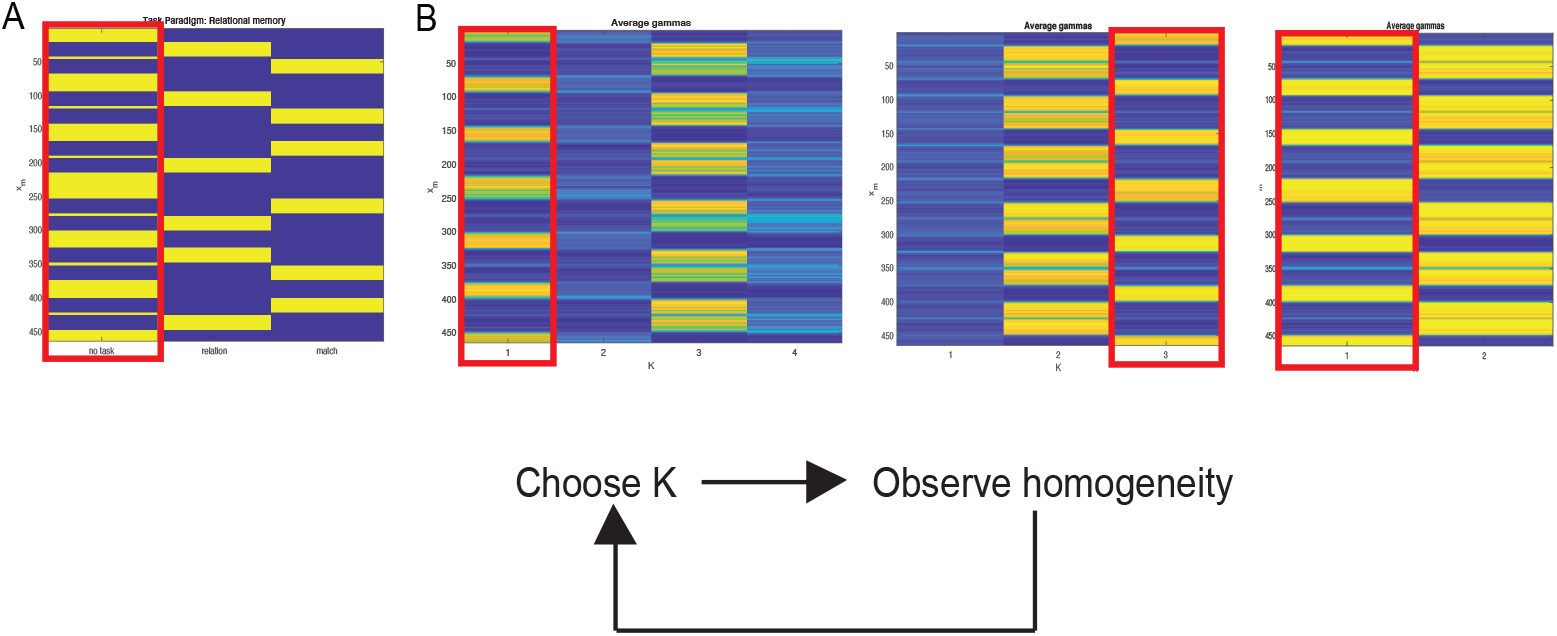
**K-***γ* **itero-homogeneity** (A) Experimental paradigm of Relational Memory task. (B)***γ*** estimated for a varying number of K sates. *K-γ itero-homogeneity* gives a visual feedback of the captured dynamics. Highlighted in red the resting states of this specific task.

#### Appendix A.4. Default Mode Networks

The DMN is the activation pattern that has been consistently found in all tasks and rest. They appear to be slightly different in their activation patterns. Their similarities in terms of connectivity and state matrices have been investigated in 3.4.

#### Appendix A.5. List of subject IDs used

As follows, the list of the 50 subjects IDs of interest from HCP: 105620, 109830, 110007, 110613, 115825, 114621, 112314, 115017, 114217, 117930, 149236, 169949, 202719, 212015, 281135, 305830, 334635, 481951, 140117, 168947, 180129, 192035, 198047, 200008, 131823, 146533, 268749, 453441, 559457, 668361, 154835, 200109, 209329, 333330, 536647, 555651, 559053, 205119, 113316, 118831, 147636, 189652, 194443, 206525, 161327, 856968, 108020, 585256, 100408, 105014, 105115, 106016, 110411, 111312, 111716, 113619, 115320, 117122, 118528, 120111, 122620, 123117, 123925, 125525, 126325, 127933, 128632, 129028, 133019, 135932, 136833, 139637, 140925, 147737, 148335, 148840, 149337, 149741, 151627, 154734, 156637, 160123, 162733, 163129, 176542, 178950, 188347, 189450, 192540, 198451, 102513, 199655, 201111, 208226, 211417, 212318, 214423, 221319, 397760, 414229, 672756, 792564, 856766, 857263, 899885.

**Figure A3:**
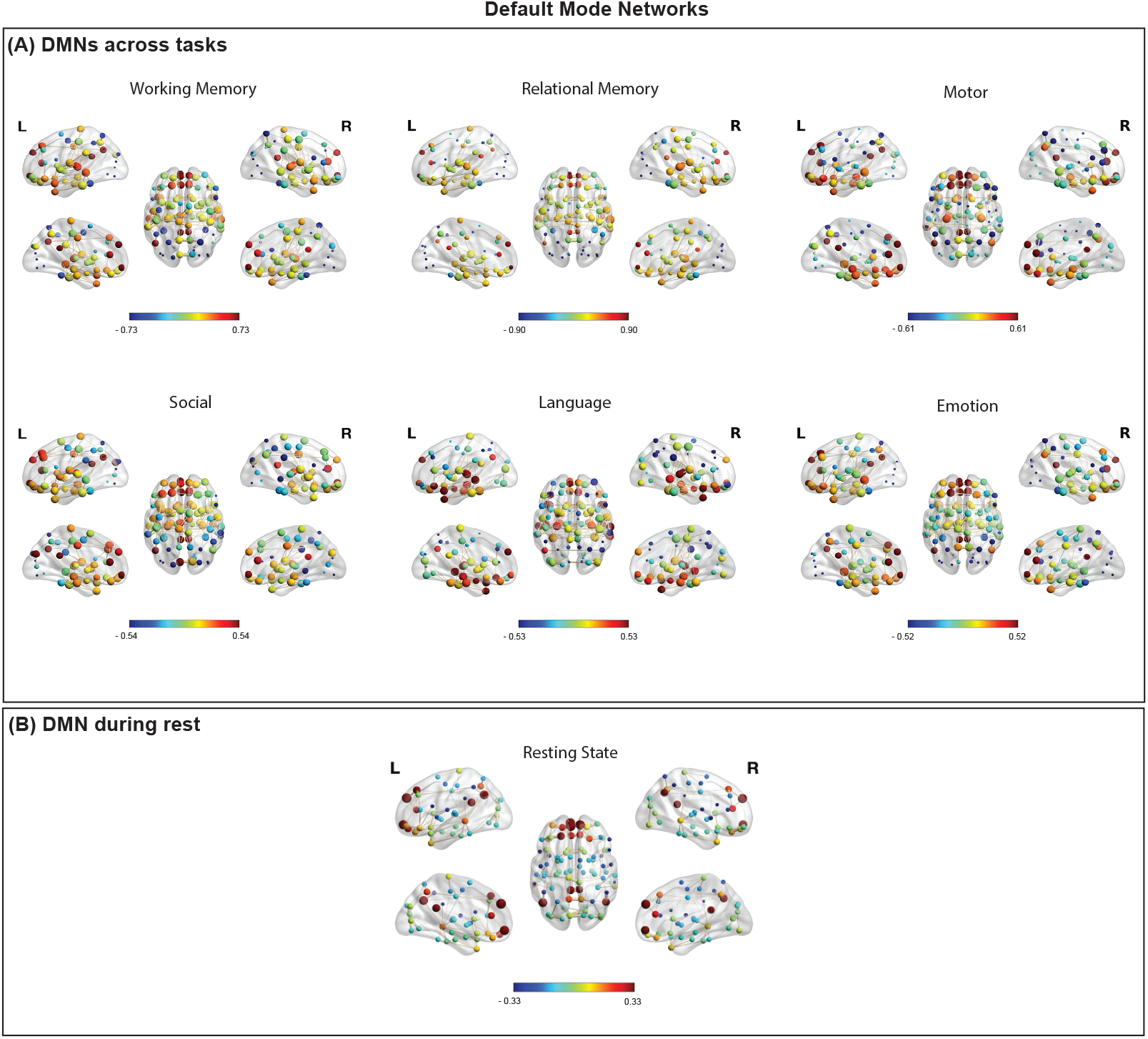
**Collection of all DMNs** (A) Default Mode Networks estimated across tasks, coinciding with the resting phase of the experimental paradigm. (B) DMN estimated from simply Resting State.

## Footnotes

1 Note that 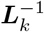 here denotes a pseudo-inverse of the graph Laplacian ***L****_k_*

2 [https://neurosynth.org/]

